# Integrative metabolomics and proteomics allow the global intracellular characterization of *Bacillus subtilis* cells and spores

**DOI:** 10.1101/2023.06.19.545067

**Authors:** Yixuan Huang, Bhagyashree N. Swarge, Winfried Roseboom, Jurre D. Bleeker, Stanley Brul, Peter Setlow, Gertjan Kramer

## Abstract

Reliable and comprehensive multi-omics analysis is essential for researchers to understand and explore complex biological systems more completely. *Bacillus subtilis* is a model organism for Gram-positive spore-forming bacteria, and in-depth insight into the physiology and molecular basis of spore formation and germination in this organism requires advanced multilayer molecular datasets generated from the same sample. In this study, we evaluated two monophasic methods for polar and nonpolar compound extraction (acetonitrile/methanol/water; isopropanol/water, and 60% ethanol), and two biphasic methods (chloroform/methanol/water, and methyl tert-butyl ether/methanol/water) on coefficients of variation of analytes, identified metabolite composition and the quality of proteomics profiles. The analytical workflow comprises (1) parallel metabolite and protein extraction, (2) monophasic or biphasic sample extraction, (3) proteomics comparison and (4) multi-omics-based data visualization. The 60% EtOH protocol proved to be the easiest in sample processing and more amenable to automation. Collectively, we annotated 505 and 484 metabolites and identified 1665 and 1562 proteins in *B. subtilis* vegetative cells and spores, respectively. We also show differences between vegetative cells and spores from a multi-omics perspective and demonstrated that an integrative multi-omics analysis can be implemented from one sample using the 60% EtOH protocol. Correlation analysis further demonstrated that the metabolome and proteome datasets were highly correlated in content for components annotated to be part of the pyrimidine metabolism pathway. The results obtained by the 60% EtOH protocol provide a comprehensive insight into differences in the metabolic and protein makeup of *B. subtilis* vegetative cells and spores.

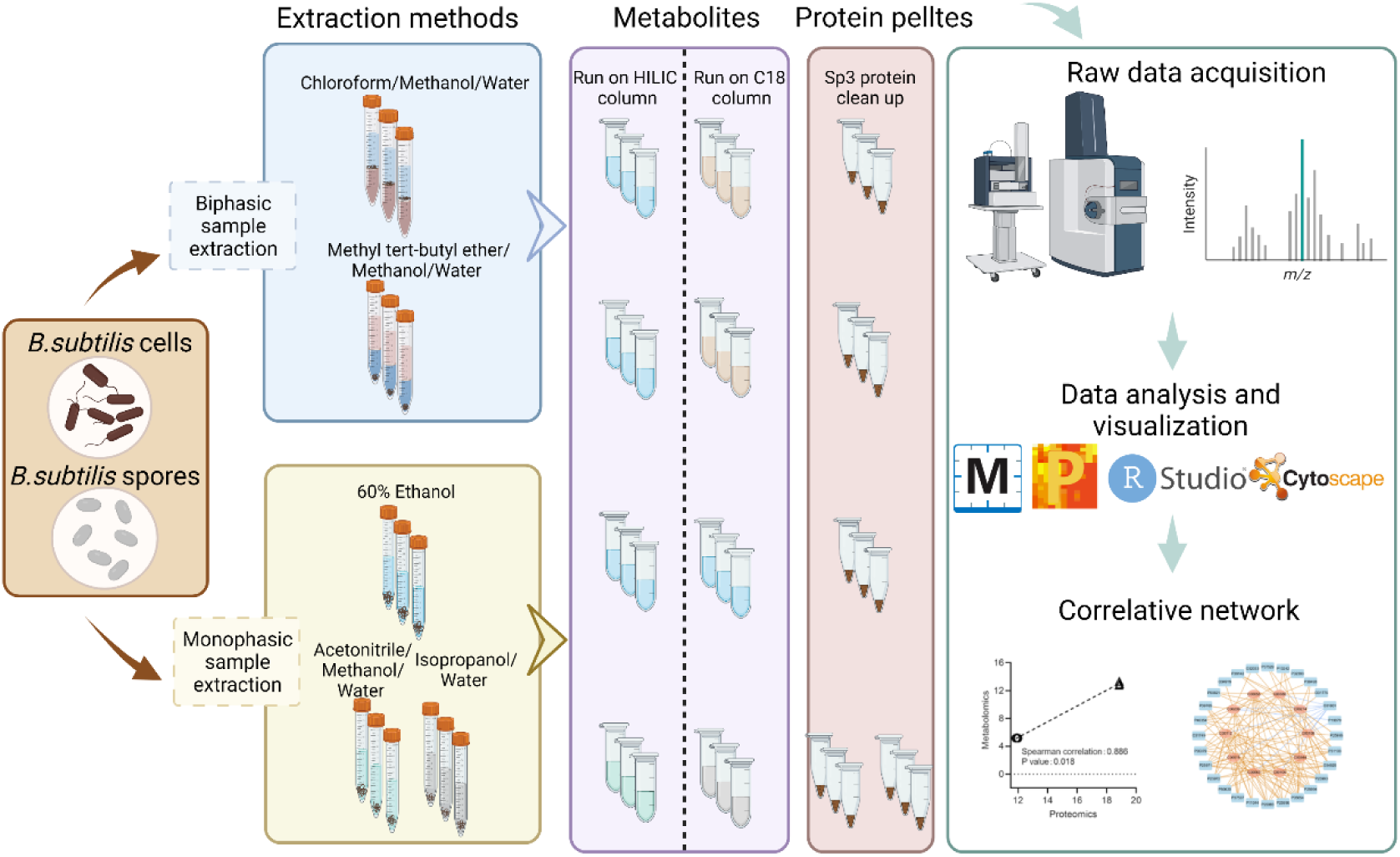

## Introduction

*Bacillus subtilis* is a prevalent Gram-positive bacterium responsible for a wide range of food spoilage in the food industry [1]. As an endospore-forming bacterium, *B. subtilis* is found in many adverse environments, such as soil, the gut of terrestrial and aquatic animals including mammals, industrial installations and healthcare facilities [2]. Survival within these extreme conditions is accomplished by integration of adaptive responses at the protein and metabolite levels leading to generation of spores with their unique multi-layered structural [3] and resistance features [4]. *B. subtilis* is a go-to model species in work to elucidate the fundamental principles of spore-formation [5]. However, while *B. subtilis* sporulation has been studied for many years, it is only recently that researchers have combined multi-level high-throughput data to understand spores’ survival mechanisms and adaptation to extreme environments at the molecular level [6,7].

With the advance of high-throughput technologies, large-scale molecular omics datasets have been widely utilized in biological research and have made possible cost-efficient, high-throughput analysis of biologic molecules [8]. The combination of multi-omics allows researchers to understand the functional changes of organisms caused by genes or environments through systematic biological information flow (DNA, RNA, protein, metabolite, and lipid), rather than just using a single omics content to find the reason for the change [9]. Recently, genomics and transcriptomics research has made tremendous progress, but this information alone cannot predict or deduce essential information at the protein, metabolite, or lipid levels [10]. For instance, changes in gene expression levels do not always correlate with translated protein abundance due to temporal shifts in the transcriptome and proteome [11]. However, in multi-omics studies that include transcriptomics or proteomics, or both, there are very few studies that also examine the metabolome [12]. Importantly, metabolomics is a powerful tool for deciphering microbial metabolism and bridging the phenotype-genotype gap because it amplifies proteomic changes and provides better characterization of biological phenotypes than any other method [13].

Multi-omics experiments often contain many different steps, each of which strongly affects the results obtained. As the first step of multi-omics research, sample preparation becomes particularly important, as the collection of reliable and highly reproducible data is crucial for drawing correct conclusions later. Due to the chemical diversity and complexity of biological components in an organism, the efficiency of the applied extraction method greatly affects the omics data, even apart from sample quality [14]. Additionally, amounts of samples are often limited, which means that it is desirable to obtain as much molecular information from a single sample as possible. Use of the same samples for multi-omics analyses will also increase consistency and comparability and decrease effort inherent in different parallel sample handling [15]. Multi-omics studies often use methods originally developed to extract either metabolites or lipids, where protein precipitation also occurs. For instance, the Folch [16] and Matyash [17] methods are biphasic extraction methods in which samples are homogenized with two immiscible lipophilic and hydrophilic solvents to simultaneously extract polar and nonpolar compounds, separating them into two solvent layers. The aqueous phase contains hydrophilic metabolites, the organic phase contains lipids and other hydrophobic metabolites, while protein is precipitated in the interphase or at the bottom of the sample. Monophasic extracts isolate water-soluble metabolites or lipids preferentially by using polar solvent(s) or nonpolar solvent(s), respectively. The biphasic methods allow extraction of polar and lipid metabolites from the same sample unlike monophasic extraction. However, monophasic protocols avoid the need to remove the biphasic liquid layers, which is difficult to automate and likely increases technical variation [18].

Recent examples of use of biphasic extraction methods in pursuit of multi-omics analysis is the use of chloroform/methanol/water (CHCl_3_/MeOH/H_2_O) to extract 1967 metabolites, 424 lipids and 1849 proteins, from a single Arabidopsis sample [19], while SIMPLEX extraction used methyl-tert-butyl-ether/methanol/water (MTBE/MeOH/H_2_O) to identify 75 metabolites, 360 lipids and 3327 proteins in mesenchymal stem cells [20]. However, simultaneous extraction of DNA (genomics), RNA (transcriptomics), proteins (proteomics), metabolites (metabolomics) and lipids (lipidomics) from a single biological sample is rare. This may be caused by the fact that optimal buffers, solutions, and protocols for extracting these very different molecules may be mutually exclusive, and longer extraction protocols can lead to molecular degradation of more labile molecular species.

In microbial metabolomics of Gram-positive bacteria such as *B. subtilis* sampling accuracy of the metabolome is key, as the highly dynamic nature necessitates fast sampling and quenching to preserve the metabolic state. The energy status of the *B. subtilis* vegetative cells, characterized by its adenylate energy charge, commonly 0.8-0.85 in actively growing cells, can be used for comparing extraction conditions and downstream analytic methods [21]. These authors found that 60% cold EtOH extraction of rapidly cooled *B. subtilis* cells isolated by vacuum filtration followed by cold water extraction gave an EC value of 0.81 ± 0.03. However, the energy charge in spores of several *Bacillus* species is ≤ 0.2 with little if any ATP present in these dormant and thought of as metabolically inactive bacterial survival structures [22,23]. Metabolic and lipidomic profiling of *Bacillus* vegetative cells have been reported [21,24,25] . However, very few studies have addressed the metabolic composition of bacterial spores, and those published focused on subsets of metabolites [22,23,26–30]. No studies to date globally compared the spore and vegetative cellular composition utilizing mass spectrometry-based metabolomics in conjunction with proteomics for multi-omics analyses. Therefore, in this study, we compare several biphasic and monophasic extraction protocols to find a reproducible and robust methodology to obtain simultaneous intracellular metabolome and proteome data profiles for *B. subtilis* spore and vegetative cell integrative multi-omics analysis. We evaluated two biphasic methods (CHCl_3_/MeOH/H_2_O, MTBE/MeOH/H_2_O), and two monophasic extraction methods i.e., 60% ethanol (EtOH) and the combination of separate extractions using acetonitrile/methanol/water (ACN/MeOH/H_2_O) and isopropanol/water (IPA/H_2_O) for polar and apolar compound extraction. The quality and repeatability as well as quantity and identity of metabolites extracted by different extraction methods were investigated by profiling intracellular metabolites with LC-MS. Furthermore, we compared the proteome coverage of vegetative cells and spore samples to control samples prepared by standard proteomics sample preparation procedures. Together, these results provide a basis for the detailed multi-omics profiling of *B. subtilis* spores and growing cells, which may lead to a deeper understanding of molecular mechanisms of spore formation and their unparalleled stress resistance properties. While our method is generally comprehensive and robust we acknowledge that improvements can still be made as, compared to classical boiling propanol extraction and in contrast to expectation, 3-PGA and nucleoside triphosphates were not robustly detected in spores and cells respectively.

## Materials and methods

### Strains and Reagents

The *B. subtilis* wild-type strain PY79 was used for this work. Tryptone, sodium chloride and yeast extract for LB (Luria-Bertani) medium were from Duchefa Biochemie (Haarlem, Netherlands). 3-(Nmorpholino) propanesulfonic acid (MOPS) used for buffered sporulation medium was purchased from Sigma Aldrich (Steinheim, Germany). Metabolite extraction solvents: Water, EtOH, MTBE, CHCl_3_ MeOH, ACN and IPA were HPLC grade and purchased from Biosolve. Ammonium bicarbonate (ABC), Sodium-dodecyl sulfate (SDS), Tris(2-carboxyethyl) phosphine hydrochloride (TCEP) and 2-chloroacetamide (CAA) used for proteomics analysis were from Sigma Aldrich. The Bicinchoninic acid (BCA) assay kit was purchased from Thermo Scientific (Schwerte, Germany), and trypsin was from Promega (Mannheim, Germany).

### Vegetative cell culturing and sporulation

A *B. subtilis* PY79 single colony was picked from an LB plate [31], inoculated in 3-5 ml LB medium (pH 7.5) and grown at 37° C in shaking incubator at 200 rpm until early log phase (OD_600_ 0.3-0.4). Serial dilutions of the log-phase cells were made in 5 ml defined sporulation minimal medium, described previously [32] and incubated at 37° C overnight. The culture with an OD_600_ of 0.3-0.4 was diluted with the pre-warmed MOPS medium and grown at 37° C in a 500 ml flask shaking at 200 rpm for 3 days, and spores harvested by centrifugation. The harvested spores were washed 3 times with chilled milliQ-water to reduce remaining vegetative cells, and Histodenz gradient centrifugation [28] was used to remove vegetative cells and phase dark cells. Only samples with more than 95% of phase bright spores as examined under a microscope were used for further research. For vegetative cell preparation, cells were harvested from LB medium upon reaching mid-exponential phase.

### Experimental design

There are 4 metabolite extraction methods used in this study, the monophasic 60% (w/v) EtOH [33], a combination of two monophasic extractions (TMEC) [34] and the bi-phasic MTBE [15] and Bligh & Dyer [35] extractions. For each method, there were three biological replicates each analyzing samples of OD_600_ = 20.

### Extraction methods

#### (i) Monophasic method - 60% (w/v) EtOH extraction

For metabolite and protein extraction and cell disruption, the harvested vegetative cell and spore pellets were transferred into 15 ml Falcon tubes containing 1 ml of cold 60% EtOH and quenched in liquid nitrogen. The whole extraction process must be carried out on ice or at 4 degrees. Subsequently, samples were thawed on ice and split into 3 bead beating tubes containing 0.5 ml 0.1 mm zirconium-silica beads (BioSpec Products, Bartlesville, OK, USA). Samples were disrupted in seven (1 min) cycles using a bacterial spore program for the OMNI bead mill homogenizer (OMNI international, Kennesaw GA, USA), which has been applied for disrupting spore and vegetative cell successfully in our previous studies [36–38]. And bead-beating has been demonstrated to yield superior extraction results compared to heat-based methods using detergent-based buffers for bacterial samples [39,40]. The samples extracted from each replicate were combined in a 15 ml Falcon tube following cell disruption, the zirconium-silica beads thoroughly washed 2 times with 1ml cold 60% EtOH to collect metabolites and cell debris from the beads. The washing solutions were combined with the metabolite extracts and centrifuged for 5 min at 4°C and 8000 rpm to harvest the supernatant for metabolomics analysis and cell debris and precipitated proteins for proteomics analysis, and both were dried under nitrogen flow. The resulting samples can be stored at -80 until analysis.

#### (ii) Monophasic method -Two monophasic extraction combination (TMEC)

In monophasic TMEC extraction, polar metabolites (ACN/MeOH/H_2_O, 1.5:1.5:1) and apolar metabolites (IPA/H_2_O, 3:1) were extracted from 10 OD at 600 nm (OD_600_) samples in two separate bead beating tubes, separately. The procedure was performed using the same steps as the 60% (w/v) EtOH extraction except for the difference in the organic solvents.

#### (iii) Biphasic method-extraction with MTBE/MeOH/H_2_O

20 OD_600_ units of vegetative cells and spores were suspended in 1 ml of cold MeOH separately and vortexed well for 30 sec. Afterwards, the quenching, bead beating and washing step were as described in the 60% EtOH extraction method, except for the different organic solvent. Then, 10 ml of MTBE was added, and samples were incubated on an orbital shaker at 100 rpm for 45 min at 4°C. To induce phase separation, 2.5 ml of water was added to each sample tube and then vortexed vigorously. Samples were centrifuged at 10000 x g for 5 min at 4 °C, and the protein pellet was in the bottom of the tube. The upper phase, lipid-containing and lower phase (polar and semi-polar metabolites) were transferred to a glass tube, and its subsequent evaporation in a speed-vacuum concentrator or nitrogen evaporator. After the supernatant was removed, the protein pellet was dried by evaporation and stored at -80°C until analysis.

#### (iv) Biphasic method-extraction with CHCl3/MeOH/H_2_O

This extraction method was performed using the same procedure as the MTBE/MeOH/H_2_O extraction, with the difference that the MTBE was replaced by CHCl_3_ and a ratio of CHCl_3_/MeOH/H_2_O of (2:2:1.8 v/v/v). Subsequently the upper polar metabolites, the lower apolar metabolites and the interphases containing protein precipitate were collected. Both different phases and pellet were dried in a centrifugal vacuum evaporator, or a nitrogen evaporator and stored at -80°C.

### Extraction of metabolites from dormant spores using boiling propanol

To evaluate whether the bead milling procedure to disrupt spores and subsequent extraction using 60% EtOH had any effect on the composition of the metabolome extracted we also extracted 20 OD_600_ units of spores as described previously [22] through direct extraction in boiling 100% 1-propanol for 5 min without mechanical disruption of spores. Extracts were lyophilized and then dissolved with cold water. Afterward, the fluids were centrifuged at 14,000 × g for 1 min, and all supernatant fractions were stored at -80°C for analysis.

### Single Tube Solid Phase Sample Preparation (SP3)

The vegetative cell or spore protein pellets obtained above were dissolved in 300 μl 1% SDS in 100 mM ABC and vortexed thoroughly. The BCA assay was used to determine the concentration of protein according to the manual and TCEP and CAA added to 10 mM and 30 mM, respectively, and the mix incubated for 0.5 h at room temperature. Samples were processed using the SP3 protein clean up [41], and trypsin (protease/protein, 1:50, w/w) was added and protein was digested at 37°C overnight. The supernatant was acidified to decouple peptides from the beads with formic acid (FA) (1% final concentration and a pH ∼2), centrifuged at 3000 g at room temperature and the supernatant moved to a clean tube, the centrifugation repeated, and the supernatant collected for LC-MS analysis.

### LC-MS/MS analysis

#### (i) Metabolomics

The dry apolar metabolites obtained by extractions by monophasic methods were reconstituted in 200 μL water while by biphasic methods were reconstituted in 200 μL IPA/ACN/water (4:3:1) and all the dry polar metabolites were in 200 μL ACN/Water (1:1). 10-μl was injected for non-targeted metabolomics onto a CSH-C18 column (100 mm × 2.1 mm,1.7 μm particle size, Waters, Massachusetts, USA) for apolar metabolites analysis or a BEH-Amide column (100 mm × 2.1 mm,1.7 μm particle size, Waters, Massachusetts, USA) for polar metabolites by an Ultimate 3000 UHPLC system (Thermo Scientific, Dreieich Germany). Using a binary solvent system (A: 0.1% FA in water, B: 0.1% FA in ACN) metabolites were separated on the CSH-C18 column by applying a linear gradient from 1% to 99% B in 18 min at a flow rate of 0.4 ml/min, while metabolites were separated on the BEH-Amide column by applying a linear gradient from 99% B to 40% B in 6 min and then to 4% B in 2 min at a flow rate of 0.4 ml/min. Eluting analytes were electrosprayed into a hybrid trapped-ion-mobility-spectroscopy quadrupole time of flight mass spectrometer (tims TOF Pro, Bruker, Bremen Germany), using a capillary voltage of 4500 volt in positive mode and 3500 volt in negative mode, with source settings as follows: end plate offset 500 volt, dry temp 250 °C, dry gas 8 l/min and nebulizer set at 3 bar both using nitrogen gas. Mass spectra were recorded using a data dependent acquisition approach in the range from m/z 20-1300 for polar and 100-1350 for the apolar metabolites in positive and negative ion mode using nitrogen as collision gas. Auto MS/MS settings were as follows: Quadupole Ion Energy 5 eV, Quadrupole Low mass 60 m/z, Collision Energy 7 eV. Active exclusion was enabled for 0.2 min, reconsidering precursors if ratio current/previous intensity > 2.

#### (ii) Proteomics

Peptides were dissolved in 6 μL water containing 0.1% FA and 3% ACN and then 200ng/μL (measured by a NanoDrop at a wavelength of 215 nm) of the peptide was injected by an Ultimate 3000 RSLCnano UHPLC system (Thermo Scientific, Germeringen, Germany). Following injection, the peptides were loaded onto a 75um x 250 mm analytical column (C18, 1.6 μm particle size, Aurora, Ionopticks, Australia) kept at 50°C and flow rate of 400 nl/min at 3% solvent B for 1 min (solvent A: 0.1% FA, solvent B: 0.1% FA in ACN). Subsequently, a stepwise gradient of 2% solvent B at 5 min, followed by 17% solvent B at 24 min, 25% solvent B at 29 min, 34% solvent B at 42 min, 99% solvent B at 33 min held until 40 min returning to initial conditions at 40.1 min equilibrating until 58 min. Eluting peptides were sprayed by the emitter coupled to the column into a captive spray source (Bruker, Bremen Germany) which was coupled to a TIMS-TOF Pro mass spectrometer. The TIMS-TOF was operated in PASEF mode of acquisition for standard proteomics. In PASEF mode, the quad isolation width was 2 Th at 700 m/z and 3 Th at 800 m/z, and the values for collision energy were set from 20-59 eV over the TIMS scan range. Precursor ions in an m/z range between 100 and 1700 with a TIMS range of 0.6 and 1.6 Vs/cm^2^ were selected for fragmentation. 10 PASEF MS/MS scans were triggered with a total cycle time of 1.16 sec, with target intensity 2e^4^ and intensity threshold of 2.5e^3^ and a charge state range of 0-5. Active exclusion was enabled for 0.4 min, reconsidering precursors if ratio current/previous intensity >4.

### LC-MS/MS Data Processing

The metabolites mass spectrometry raw files were submitted to MetaboScape 5.0 (Bruker Daltonics, Germany) used to perform data deconvolution, peak-picking, and alignment of m/z features using the TReX 3D peak extraction and alignment algorithm (EIC correlation set at 0.8). All spectra were recalibrated on an internal lockmass segment (NaFormate clusters) and peaks were extracted with a minimum peak length of 8 spectra (7 for recursive extraction) and an intensity threshold of 1000 counts for peak detection. In negative mode ion deconvolution setting, [M-H]-was set for primary ion, seed ion was [M+Cl]- and common ions were [M-H-H_2_O]-, [M+COOH]-. For positive mode, the primary ion was [M+H] +, seed ions were [M+Na] +, [M+K] +, [M+NH_4_] + and [M-H-H_2_O] + was the common ion [42]. Features were annotated, using SMARTFORMULA (narrow threshold, 3.0 mDa, mSigma:15; wide threshold 5.0 mDa, mSigma:30), to calculate a molecular formula. Spectral libraries including Bruker MetaboBASE 3.0, Bruker HDBM 2.0, MetaboBASE 2.0 in silico, MSDIAL LipidDBs, MoNA VF NPL QTOF, AND GNPS export, were used for feature annotation (narrow threshold, 2.0 mDa, mSigma 10, msms score 900, wide threshold 5.0 mDa, mSigma:20 msms score 800). Analyte listS containing 667 compounds with a retention time (RT) (narrow threshold, 1.0 mDa, 0.05 min, mSigma: 10, msms score 900; wide threshold 5.0 mDa, 0.1 min, mSigma 50, msms score 700) was also used to annotate deconvoluted features. An annotated feature was considered of high confidence if more than two green boxes were present in the Annotation Quality column of the program and low confidence if less than two green boxes were present. Resulting data were exported for further analysis with MetaboAnalyst 5.0 (https://www.metaboanalyst.ca/MetaboAnalyst/home.xhtml). At first, the data showing a poor variation were filtered on Inter Quartile Range (IQR) and then features were normalized by median normalization, scaled by Auto scaling and transformed to a logarithmic scale (base of 2).

Generated Mass spectra for pellets (vegetative cells and spores) were analyzed with Maxquant (Version 1.6.14) for feature detection and protein identification. Tims-DDA was set in Type of Group specific and other parameters were set as the default. Searches included variable modifications of methionine oxidation, and a fixed modification of cysteine carbamidomethyl and the proteolytic enzyme was trypsin with maximum 2 missed cleavages. A *Bacillus subtilis* database (version 2019 downloaded from Uniprot) was used for database searches. To improve mass accuracy of matching precursors, the “match between runs” option was applied within a match window time of 0.2 min and a match ion mobility window of 0.05. Proteins of label free quantification (LFQ) calculated for each pellet represented normalized peptide intensities correlating with protein abundances. Finally, all the quantification and annotation information were summed in in the output proteinGroup.txt.

Perseus (1.6.15.0) was processed for analysis of Maxquant results (proteinGroup.txt). Briefly, the LFQ intensity of each sample was selected as the main data matrix and then potential contaminants, reverse and only identified by site were removed. Afterwards, the LFQ intensity of proteins of which the unique peptides >1 was transformed to log2[x], and proteins in at least two of the three replicates were further analyzed. Normalization was achieved using a Z-score with matrix access by rows (rows are proteins, columns are samples). K means was used for the hierarchical clustering of rows (proteins) and significant protein expression differences between different extractions were identified using p values< 0.05 from student’s test.

Calculation and plotting for principal component analysis (PCA) of metabolome profiles were implemented with SIMCA software. Kyoto Encyclopedia of Genes and Genomes (KEGG) enrichment analysis of differentially expressed proteins was implemented by the ClusterProfiler R package. For differentially expressed metabolites, MetaboAnalyst 5.0 (https://www.metaboanalyst.ca/<u>) was</u> applied for KEGG pathway enrichment. Gene ontology (GO) enrichment analysis was applied with geneontology (http://geneontology.org/). For metabolites and proteins correlation network calculation and visualization, R package WGCNA and Cytoscape (Version 3.8.2) were used.

## Results

### Comparison of the metabolome and lipidome extracted from cells and spores

Given that extraction methods can alter the measurable metabolome and lipidome we first set out to compare how different extraction methods affect mass spectrometry-based analysis of cells and spores of *B. subtilis*. To assess how the four extraction methods, monophasic: EtOH, TMEC and biphasic: MeOH/MTBE, MeOH/CHCl_3_, performed we assessed the number of metabolite features extracted, the variability of quantitation and compared if there is any bias in classes of detected molecules. For bacterial cells the biphasic methods led to the detection of most features (MeOH/MTBE > MeOH/CHCl_3_ > EtOH > TMEC), while in spores, monophasic methods (TMEC > EtOH > MeOH/CHCl_3_ > MeOH/MTBE) detected most features when comparing m/z features detected in all three biological replicates (Table I). Reproducibility of detection of m/z features was also best for biphasic extractions in bacterial cells with MeOH/CHCl_3_ having the lowest median CV (0.24) and TMEC (0.31) the highest. Conversely the monophasic extraction TMEC had the lowest median CV (0.15) in bacterial spores for m/z features detected in all three replicates (Table I and Fig. S1). Amongst the detected m/z features some were putatively annotated with a molecular structure (Table I). Following removal of redundancy (features being assigned the same putative structure) the four protocols shared 338 cellular metabolites of 598 total and 281 spore metabolites of 532 total (Fig. 1 A,B). Of the extraction methods the monophasic EtOH method had most putatively annotated metabolites in both vegetative cells (506) and spores (484), as well as most that were only found in this extraction method (34 in cells and 50 in spores).

**Table I.**
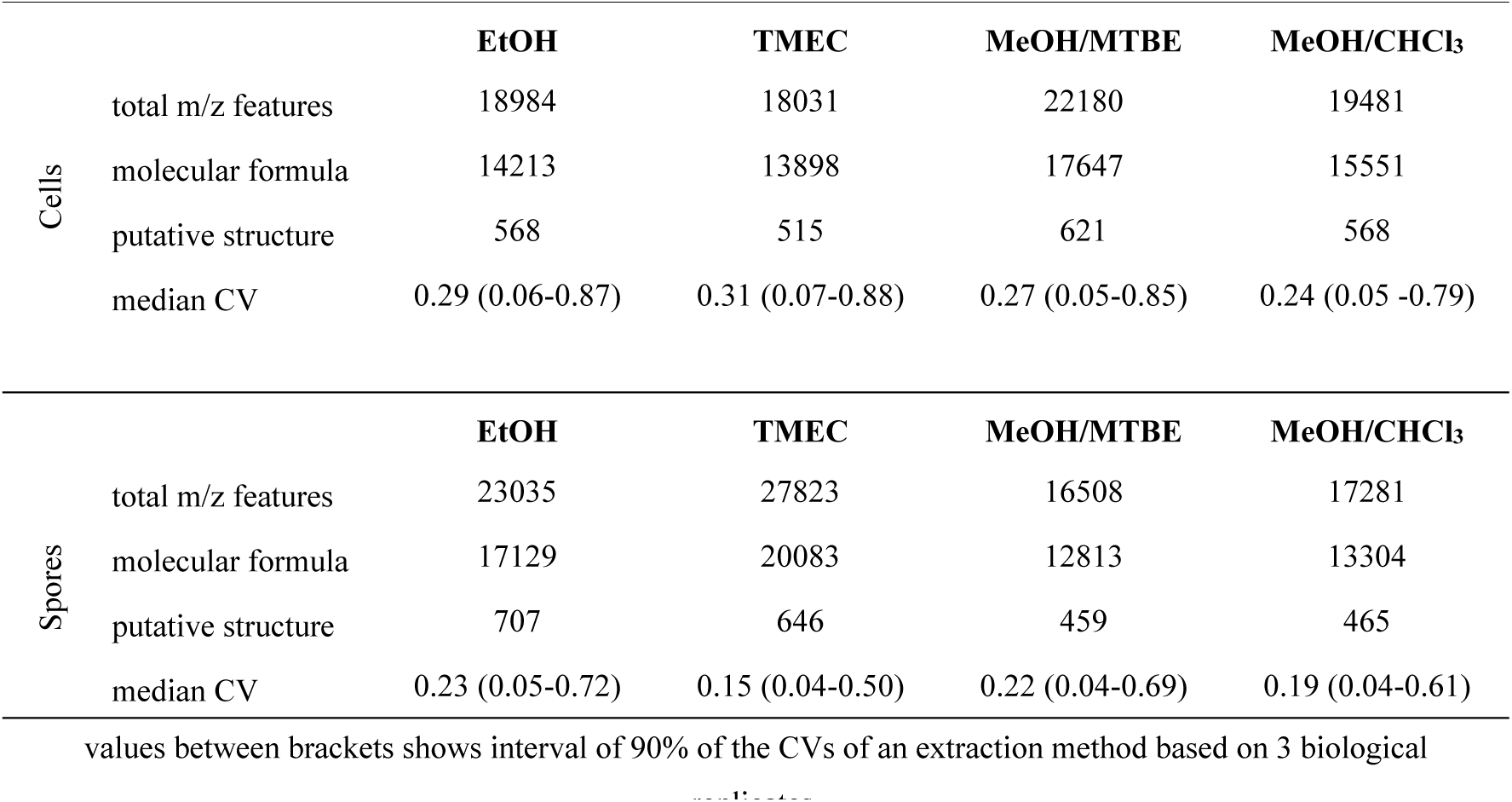
Molecular features detected using different extraction methods.

**Fig. 1.**
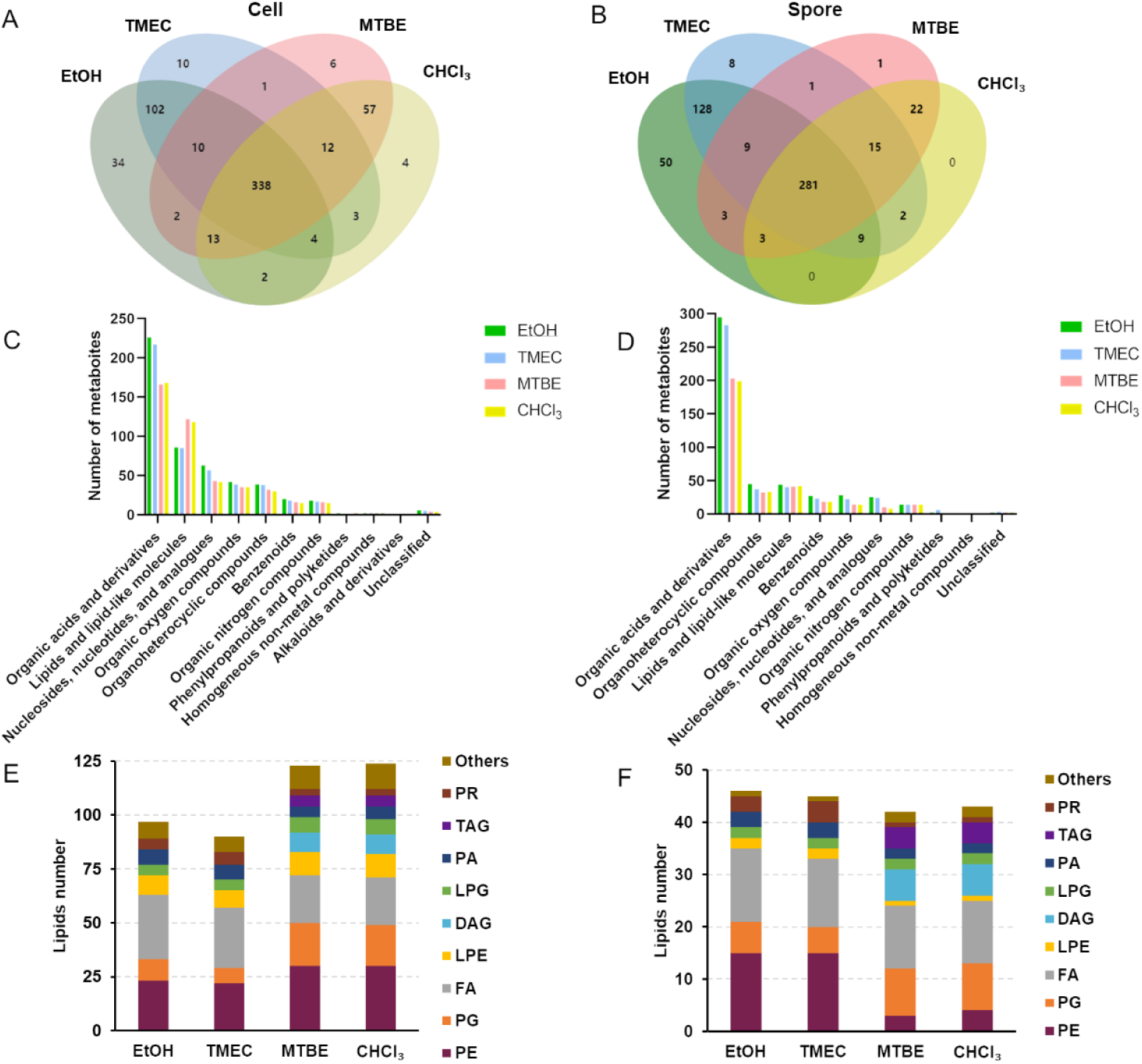
Comparison of metabolites identified by different extraction methods in *B. subtilis* cells and spores. (A and B) Venn diagram showing the overlapping (shared) and unique metabolites identified in the cells and spores in different methods. (C and D) Bar graphs of metabolites showing the difference in metabolite classes in cells and spores extracted by different methods. (E and F) Comparison of distribution of annotated lipids in major lipid classes identified by different extraction methods in *B. subtilis* cells and spores.

To assess whether there was any specific bias in the type of molecules extracted by the different methods we assigned the non-redundant putative annotations into molecular categories (Fig. 1 C,D). Strikingly both monophasic methods show a markedly higher number of molecules in the categories of organic acids and derivatives and nucleosides, nucleotides, and analogues in both spores and cells compared to the biphasic methods. On the other hand, the biphasic methods had a larger number of molecules in the category of lipids and lipid-like molecules in vegetative cells but not spores. We examined the categories of the lipids extracted by the 4 different methods from cell and spore samples more closely (Fig. 1 E,F). The biphasic methods found a larger number of lipids (124 vs. 97) in vegetative cells, while lipids detected differed little (43 vs. 46) in spores between the biphasic and monophasic methods. Strikingly diacyl- and triacyl-glycerols (DAG and TAG) were only detected in samples extracted by biphasic methods and phosphatidylethanolamines (PE) were mainly found in samples extracted by mono-phasic methods in spores (15 vs. 4) and by biphasic methods in vegetative cells (30 vs. 23). In terms of solvent polarity, MTBE and CHCl_3_ are more non-polar than EtOH and IPA, explaining why biphasic methods show a better coverage of lipid and lipid-like molecules. This was less obvious in spore samples, even though, as was shown in *Bacillus licheniformis*, the total lipids of vegetative cells (2.9% of the dry weight) is not very different of that of spores [43] (2.1% of the dry weight). We additionally compared the 60% EtOH extraction to extraction in boiling propanol which is a method that has been used previously in small molecule analysis of *Bacillus* spores [22]. We assessed whether mechanical spore disruption or residual enzyme activity had any large influence on the quantitation of tentatively annotated molecules from these spores, but no large bias was obvious, although on average nucleotide extraction seemed better with boiling propanol (Fig. S2 and Table S1). Noticeably, compared to boiling propanol extraction and in contrast to expectation, 3-PGA and nucleoside triphosphates were not well detected in spores and cells respectively.

Overall, depending on the background (spores or cells) different extraction methods seems to provide different benefits and drawbacks in terms of reproducibility, number of features detected, and types of molecules annotated without one clear-cut best approach. For multi-omics however, small molecule extraction is only one part of the performance to take into consideration, the other is protein extraction and proteome coverage.

### Proteome coverage of B. subtilis cells and spores using different extraction methods

Since the 4 extraction protocols assayed in the above were originally established for the extraction of small molecules, we next focused our efforts towards assessing their utility in extracting proteins for proteomics analysis. To establish an optimal protocol for multi-omics analyses from a single sample, we investigated if protein fractions isolated by the two monophasic and biphasic extraction procedures yield similar results compared to extracting proteins directly with 1% SDS (n = 3 for all) which is a common lysis buffer in proteomics. To do so we also dissolved the protein fractions of *B. subtilis* cells and spores in 1% SDS and used SP3 to clean up all samples prior to digestion with trypsin. Analysis of the protein fractions from the different small molecule extraction methods show similar or more proteins identified and quantified in cells or spores (detected in all three replicates) compared to direct extraction using 1% SDS (Table II). Notably most proteins identified were from the EtOH extraction in cells (1785 proteins) and from the IPA extraction of TMEC in spores (1659 proteins). Obtaining reproducible quantitative information is critical to identify differential protein expression for downstream analysis in proteomics studies. Therefore, we compared the variance in protein quantitation between the different methods, and median CV showed little difference (ranging between 0.10-0.14) between the different extraction methods compared to the standard extraction using 1% SDS (Table II and Fig. S3 A,B). As such, these methods seem on par or better when compared to the standard extraction method in proteomics when regarding these metrics.

**Table II.** Proteins detected using different extraction methods.

|  |  | **EtOH** | **TMEC** | **TMEC** | **MeOH** | **MeOH** | **1% SDS** |
| --- | --- | --- | --- | --- | --- | --- | --- |
|  |  |  | **ACN** | **IPA** | **CHCl3** | **MTBE** |  |
| Cells | protein id | 1785 | 1491 | 1645 | 1576 | 1565 | 1570 |
|  | prot. quant. | 1479 | 1120 | 1275 | 1185 | 1164 | 1199 |
|  | median CV | 0.11 | 0.11 | 0.12 | 0.13 | 0.13 | 0.13 |
|  |  | (0.02-0.37) | (0.02-0.38) | (0.03-0.37) | (0.03-0.43) | (0.03-0.40) | (0.03-0.45) |
| Spores | protein id | 1443 | 1659 | 1626 | 1562 | 1522 | 1433 |
|  | prot. quant. | 1026 | 1329 | 1306 | 1242 | 1172 | 1015 |
|  | median CV | 0.12 | 0.11 | 0.10 | 0.14 | 0.13 | 0.10 |
|  |  | (0.02-0.50) | (0.02-0.47) | (0.02-0.45) | (0.04-0.51) | (0.03-0.48) | (0.02-0.37) |

values between brackets shows interval of 90% of the CVs of an extraction method based on 3 biological replicates.

Apart from reproducible quantitation, the method of extraction can change the composition of the proteome under study, while digesting proteins using different buffers, surfactants, or denaturing agents can influence the proteome coverage [44]. The extraction reagents used in this study are all organic solvents (varying in polarity), which means their precipitating effect on proteins is mediated by increasing the attractive forces between the protein molecules and causing a dehydration effect on the proteins to facilitate the interaction between them [45,46]. The overlap of proteins identified from all different extraction methods is very high (Fig. S3 C,D) with 1231 proteins and 1157 proteins in vegetative cells and spores found irrespective of extraction method. While 92 (cells) and 33 (spores) proteins were specifically obtained by one of the metabolite extraction methods compared to 1% SDS extraction and only 4 proteins were unique to the 1% SDS extraction. Proteins quantified following the different extraction methods are shown in a heatmap to show the overall differences in representation of individual proteins in the datasets (Fig. 2 A, C). The quantified proteins are classified into 3 clusters according to K-means clustering and these clusters were also annotated by Gene Ontology terms regarding Cellular Component (GOCC). In cells, 60% EtOH was highly similar in protein quantitation as 1% SDS in cluster 1 but had lower intensities of proteins in cluster 2 which was enriched in proteins of the cell periphery and membrane and higher amounts of proteins in cluster 3 which was enriched for ribosomal proteins (Fig. 2B). There was little difference in the other 4 extraction methods. In spores, the 1% SDS extracted a larger relative quantity of proteins in clusters 1 and 3 which are enriched in the GO terms ribosomal subunit and cell wall, but less in cluster 2 (Fig. 2D). Overall, there is no significant evidence that the metabolite extraction methods affected the coverage of the proteome in our experiments indicating it is feasible to do simultaneous metabolomic and proteomic analyses from the same sample in *B. subtilis* cells and spores. Considering performance on these multiple omics and convenience in sample preparation we chose to continue with the 60% EtOH extraction to analyze the differences in the spore and cellular metabolome, lipidome and proteome of *B. subtilis* below.

**Fig. 2.**
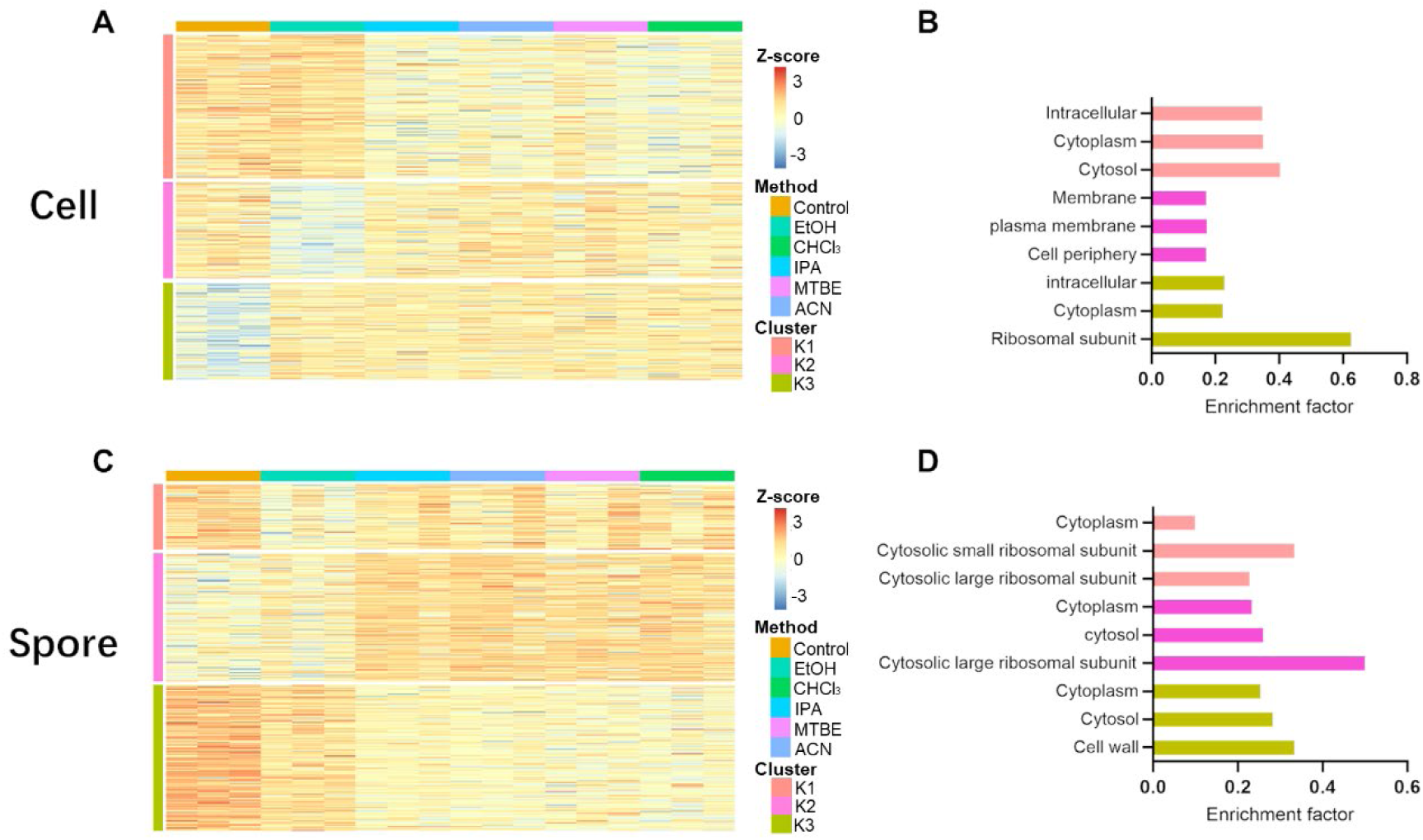
Differences in proteomic profiles of different extraction methods. (A and C) Heatmap of the quantified proteins from the different extraction methods. Proteins from cell and spores are clustered in 3 (K1, K2, K3) in K-means cluster analysis. (B and D) Bar graphs of fractions of the cluster classified to GOCC categories. Only the top 3 categories are shown in the graphs. Colors of the bars correspond to the clusters in the heatmaps.

### The metabolome of vegetative cells and spores

Many studies have examined the properties of *B. subtilis* that show significant changes in all kinds of resistance with the change of physiological form [47]. However, no studies have tried to globally compare the metabolite content between vegetative cells and spores to obtain a complete view of the molecular composition of these two distinct states of the bacterial life cycle. Comparison of the m/z features that had a putative annotation of a molecular structure showed that 213 metabolites were detected in both datasets, with 292 metabolites uniquely found in cells and 271 metabolites only found in spores (Fig. 3A) Metabolites that were annotated in vegetative cells but not in spores or vice versa are likely predominant in cells and spores and potentially important, we further referred to these as cell or spore specific metabolites. However, differences in dynamic range of the metabolome between these two distinct forms of *B. subtilis* can also lead to non-detection of some of these metabolites in one of the two sample types. Amongst the spore specific molecules with a putative annotation was dipicolinic acid which is a well-known and abundant molecule involved in spore resistance to a variety of stresses. Next, we examined the differential expression of these 213 shared metabolites by performing a pair wise comparison. Log2 fold changes greater than 1 (or less than -1) and -Log10 P value greater than 1.31 indicate metabolites that were predominant in cells (red) or spores (blue, Fig. 3B). A total of 117 metabolites were differentially present in vegetative cells or spores. We performed KEGG pathway enrichment analysis with all the differentially present metabolites and specific metabolites. Enrichment analysis showed that the altered molecules belonged to 15 pathways including arginine biosynthesis and purine metabolism (Fig. 3C). The detailed relative levels of these metabolites in vegetative cells and spores of these KEGG pathways are shown in Fig. 3D. These data show that some small molecules exhibit a significantly higher accumulation in spores compared to bacteria, with active metabolism, presumably stored during sporulation.

**Fig. 3.**
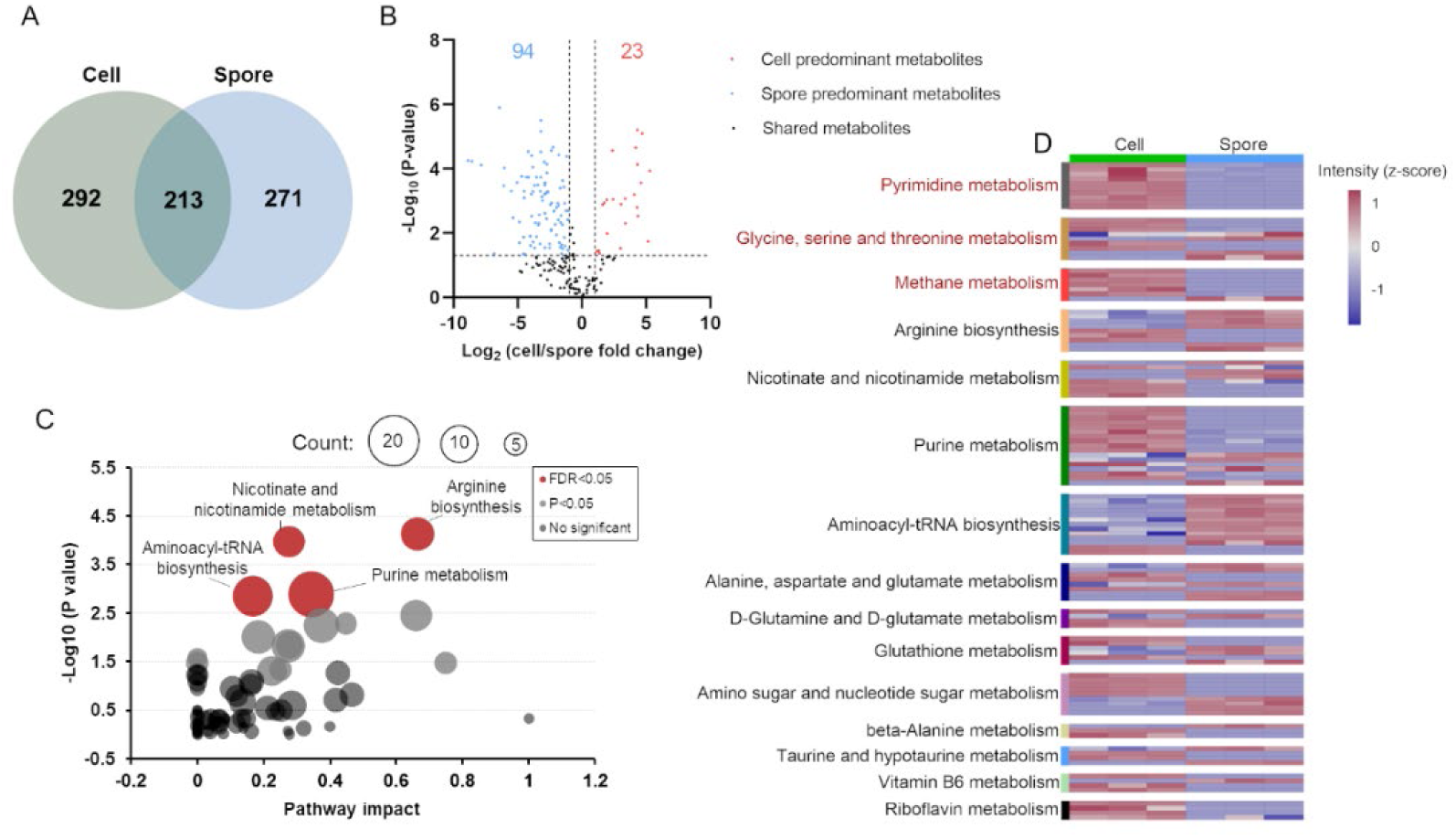
Analysis of metabolome profiles of *B. subtilis* vegetative cells and spores extracted by 60% EtOH. (A) A Venn-diagram illustrating the total numbers of annotated compounds uniquely found in cells and spores and the shared metabolites. (B) Volcano plot of comparison between cell and spore metabolomes. Student t test (P< 0.05) and Fold changes>1 or <-1 were in red or blue dots. (C) The KEGG pathway enrichment analysis of significant metabolites. The *x*-axis represents pathway impact, and the *y*-axis represents the −log10(p value). The dot size represents the metabolite numbers in the pathway. The red color indicates the enriched pathways (FDR< 0.05). (D) A heatmap showing the expression levels of 15 enriched metabolic pathways in cells and spores. The color legend indicates the *z*-score transformed intensity of the pathway.

### Analysis of the cell and spore proteome of B. subtilis

To further explore the differences between vegetative cells and spores, we quantitatively investigated the proteomes by analysis of proteins precipitated during EtOH extraction. Following analysis, 1665 proteins were identified and quantified in vegetative cells and 1562 in spores, which is a distinct improvement over our prior study of *B. subtilis* spores and cells where we quantified 1086 spore and cellular proteins [7]. Following stringent filtering 568 proteins were found to be specific for vegetative cells and 465 were uniquely identified in spores (Fig. 4A). Amongst the uniquely identified in spores unsurprisingly many structural spore and spore coat proteins were found as well as proteins related to spore revival, while the proteins detected in vegetative cells were related to cellular mobility amongst others. Of the 1097 proteins detected in both stages of the lifecycle of *B. subtilis*, 693 were predominant in vegetative cells (log2 fold change >1, p-value < 0.05) while 94 proteins were predominant in spores (log2 fold change <-1, p-value < 0.05, Fig. 4B). These life cycle specific and predominant proteins were searched for functional enrichment of GO-terms to obtain an overview of molecular function (MF), cellular component (CC), and biological process (BP) differences when *Bacilli* transition from vegetative cells to spores (Fig. 4D). This identified the leading terms as mostly associated with cellular metabolic processes and cellular components. To complement these analyses, we also looked for enrichment in the KEGG database to explore the unique and common functional pathways. The KEGG enriched data were classified into three groups (i.e. FDR< 0.05, P-value< 0.05, not significant) from which the top 10 pathways, (red dots Fig. 4C) were used for further analysis. Starting from these significantly enriched pathways, a heatmap was constructed to show the expression level in vegetative cells and spores. Results showed that, most proteins significantly enriched with the term “the biosynthesis of secondary metabolites” (Fig. 4E) had a higher expression in vegetative cells.

**Fig. 4.**
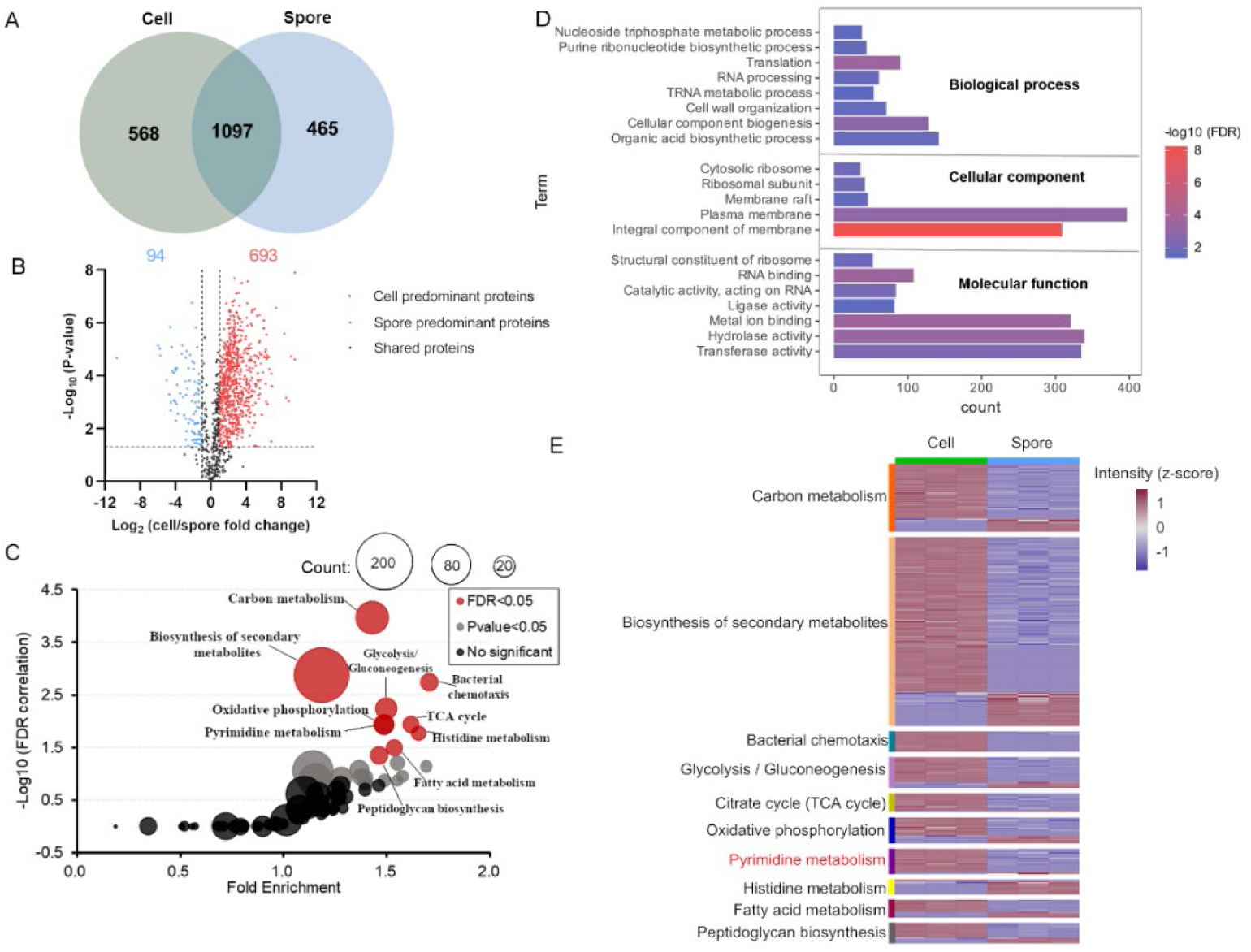
Analysis of proteomics profiles of *B. subtilis* vegetative cells and spores extracted by 60% EtOH. (A) Venn-diagram illustrating the total numbers of proteins identified uniquely or shared in cells and spores. (B) Volcano plot of comparison between the cell and spore proteomes. Student t test (P< 0.05) and Fold changes>1 or <-1 were in red or blue dots. (C) The x-axis represents fold enrichment and the y-axis represents the −log10(FDR). The dot size represents the protein numbers in the pathway. The red color indicates the enriched pathways (FDR< 0.05). (D) GO analysis of differentially and unique proteins showing the items with FDR < 0.05. (E) A heatmap showing the expression levels of 10 enriched metabolic pathways. The color legend indicates the z-score transformed intensity of proteins.

### Integral analysis of omics from vegetative cells and spores

To illustrate the value of multiple omics measurements on the same sample, we performed integrative pathway analysis of differentially expressed metabolites and proteins found in the analysis based on metabolomics and proteomics results of vegetative cells versus spores. This analysis comprised 3 common KEGG pathways (pyrimidine metabolism, glycine, serine and threonine metabolism and methane metabolism), and a Spearman correlation between metabolomics and proteomics datasets was calculated. Only the pyrimidine metabolism pathway showed a Spearman correlation > 0.7 and p-value <0.05 between changes in metabolites and protein abundance in vegetative cells versus spores (Fig. 5A). The pyrimidine metabolism pathway was highly correlated at both the level of metabolite and proteins expression changes with correlation coefficient of 0.89 and p-value of 0.018. The correlation network shown in Fig. 5B consists of 133 edges containing 28 proteins and 10 metabolites, demonstrating a significant coherency between changes in the composition of metabolites and proteins between these two states.

**Fig. 5.**
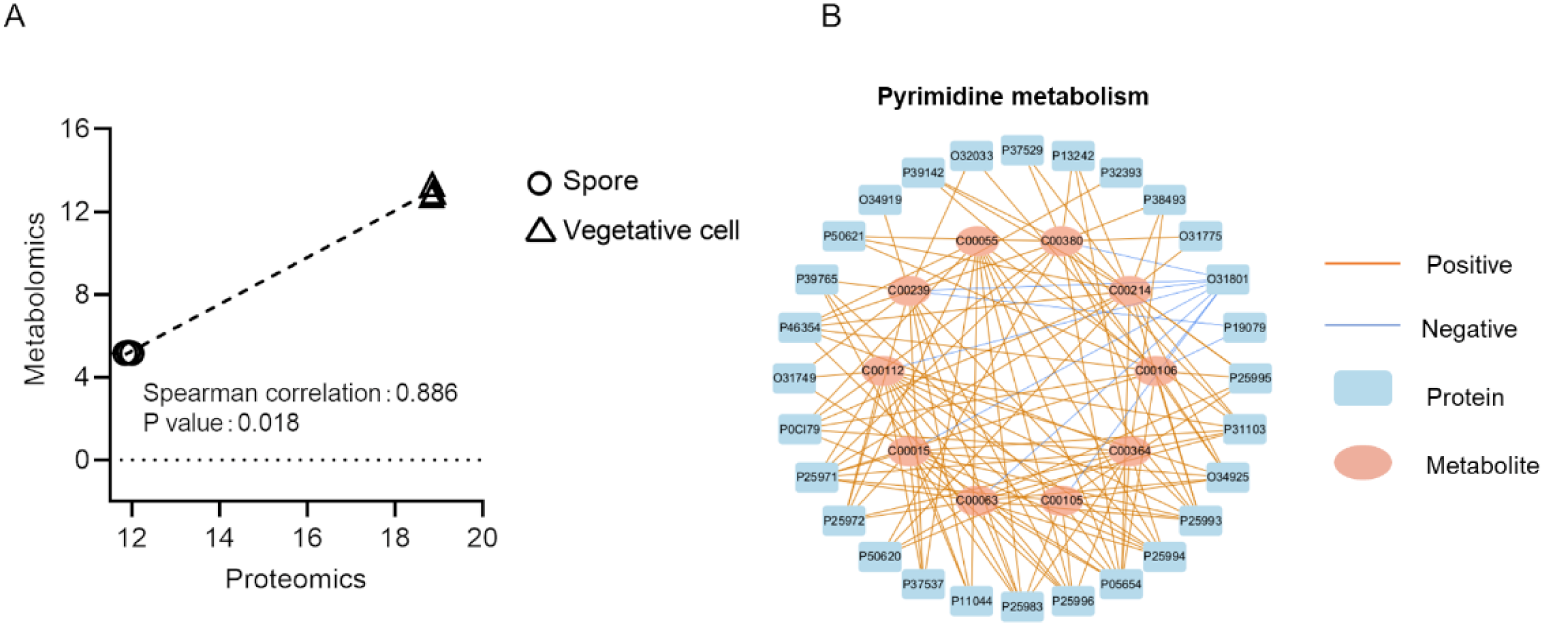
Integrative pathway enrichment analysis of multi-omics data extracted by 60% EtOH. (A) Correlation of the expressions for the pyrimidine metabolism pathway between proteomics and metabolomics. (B) The correlation network of proteins and metabolites in the pyrimidine metabolism pathway. Each node represents one protein or one metabolite. The connections were established by Spearman correlation (Student’s *t* test, *P* values < 0.05 and absolute Spearman correlation values > 0.7).

Through studying the comparison between vegetative cells and spores, the results show that 60% EtOH extraction is suitable to identify the cross talk between metabolites and proteins. The correlative effects between metabolome and proteome give additional information on the inter-molecular regulation of biological pathways, which is not possible by using only a single omics analysis.

## Discussion

Although multi-omics studies have significantly increased researchers’ potential towards understanding complex biological systems, at the same time multi-omics studies face challenges in different aspects, such as study design and variable depth of analysis of different omics and the complexity of subsequent integrative data analysis. Among the types of integrated multi-omics studies published, the combination of transcriptomics and proteomics is the most common, followed by the integration between transcriptomics and genomics [48]. Metabolomics is a young field compared to other omics in which untargeted metabolomics provides an unbiased, high-coverage metabolome, important for the discovery phase of research. However reproducibility of analysis can be problematic and it requires complex data processing [49]. Therefore, there are few studies integrating metabolomics and none to date comparing vegetative cells and spores. But for the most direct information about the physiological state or phenotype, integrating large-scale quantification of protein and metabolite content provides the best description of the biological system. Thus, finding an optimal and robust sample preparation method that can sufficiently address both the proteome and metabolome is a first essential step for robust multi-omics analysis.

This study provides a comprehensive and comparative analysis of the metabolome and proteome of *B. subtilis* vegetative cell and spore samples. We compared 4 different extraction methods regarding the composition of metabolites and proteins found as well as the reproducibility of quantification and the number of identified metabolites and proteins. We propose extraction with 60% EtOH, of which the advantages are (i) easily to operate and avoids contamination between different polarity layers, (ii) less procedures leading to low sampling errors, (iii) higher number of identified metabolites, (iv) low variation of the proteome, (v) no qualitative losses of the proteome composition, and (vi) low toxicity in line with lab safety and an environmental sustainability. In-depth classification with lipid composition indicated that the method identified less lipids than the biphasic approaches. Nonpolar solvents combined with polar solvents have the most potential to extract lipids [50,51]. The neutral lipids are dissolved by nonpolar solvents, but neutral lipids which are associated with polar lipids via hydrogen bonding can remain behind in the biological sample, a polar solvent is used to break these hydrogen bonds and extracting these lipids along with polar lipid [52]. Here the 60% EtOH extraction does extract lipids but is somewhat less suited for studies focused on lipid detection.

The application of 60% EtOH and identification of a close correlation between small molecules and protein abundance was shown by studying the composition of *B. subtilis* vegetative cells and spores. In contrast to the study of the transcriptome, proteome or both, far fewer studies have focused on combining intracellular metabolomics with proteomics in a multi-omics approach [12,53]. But, as important as it is to correlate gene and protein expression to study post-transcriptional regulation and phenotypic-function, the availability of co-factors (e.g. calcium, zinc, ATP, NAD), flux and activities of each enzyme also need to be considered to fully describe the regulation of biological processes [54]. Therefore, we attempted to elucidate the difference between *B. subtilis* vegetative cells and spores within the broader context of both protein and small molecular composition to gain a more complete overview of the interaction between metabolite and protein content. We profiled the *B. subtilis* vegetative cell and spore proteome and metabolome by 60% EtOH extraction followed by LC-MS analysis. Our data showed low technical variation, enabling insight into metabolite and protein composition within spores and cells. We found 3 common differentially abundant pathways in both metabolomics and proteomics and one prominent example was the pyrimidine metabolism pathway, which showed high coherency in differences of metabolites and protein quantities. Through the construction of the association network between metabolites and proteins, O31801 (*yncF*) and P19079 (*cdd*), which were negatively correlated with metabolites, attracted our attention. In nucleotide metabolism, deoxyuridine 5’-triphosphate pyrophosphatase (YncF) produces 2’-deoxyuridine 5-monophosphate (dUMP) and decreases the intracellular concentration of dUTP, preventing DNA uracil incorporation. Uracil, however, can originate from cytosine deamination and is one of the most frequent erroneous bases in DNA [55], and cytidine deaminase (CDA, encoded by the *cdd* gene) is responsible for deaminating cytidine to uridine.

As structural components of several key molecules, pyrimidines play an important role in a wide range of cellular functions, including DNA and RNA synthesis, as a component of triphosphates (UTP and CTP) [56]. Vegetative cells express a high level of metabolites and proteins involved in pyrimidine metabolism, which is in accordance with previous findings [7]. To complete bacterial replication, vegetative cells growing logarithmically require a considerable amount of energy and nucleotides. In contrast, there has not been any report concerning the altered expression of the *yncf* and *cdd* enzymes in spores. The transition from vegetative cells to spores reflects the differences that affect developmental programs (such as sporulation), suggesting that these pathways are plastic in nature and may be under greater environmental selection [57].

Overall, we compared multi-omics sample extraction protocols for *B. subtilis* vegetative cells and spores, which provides a window on best practices for future systemic integrative studies on sporulation. Our research concluded that 60% EtOH extraction worked easily with more identified metabolites and low variation for an integrative metabolomics and proteomics study of *B. subtilis* vegetative cells and spores. We are aware of the limitation of the 60% EtOH extraction method not resolving, in contrast to expectation, 3-PGA and nucleoside triphosphates in spores and cells respectively. Nonetheless, we are convinced that this large-scale comparative study of four different extraction approaches for the extraction provide a comprehensive and detailed dataset in *B. subtilis* multi-omics methodology studies that will be beneficial for further study on multi-molecular cellular processes during spore formation and spore revival.

## Supporting information

Data Underlying Figures in Manuscript

## Author contributions

Conceptualization, S.B., G.K. and Y.H.; methodology, Y.H., B.N.S. and J.B.; formal analysis, Y.H.; writing—original draft preparation, Y.H.; writing—review and editing, S.B., P.S., G.K. and B.N.S.; resources, W.R. All authors have read and agreed to the published version of the manuscript.

## Data Availability

All mass spectral data have been deposited into Massive (https://massive.ucsd.edu/ ProteoSAFe/static/massive.jsp), with the identifier MSV000092140

## Supplemental Figures and Tables

**Figure S1.**
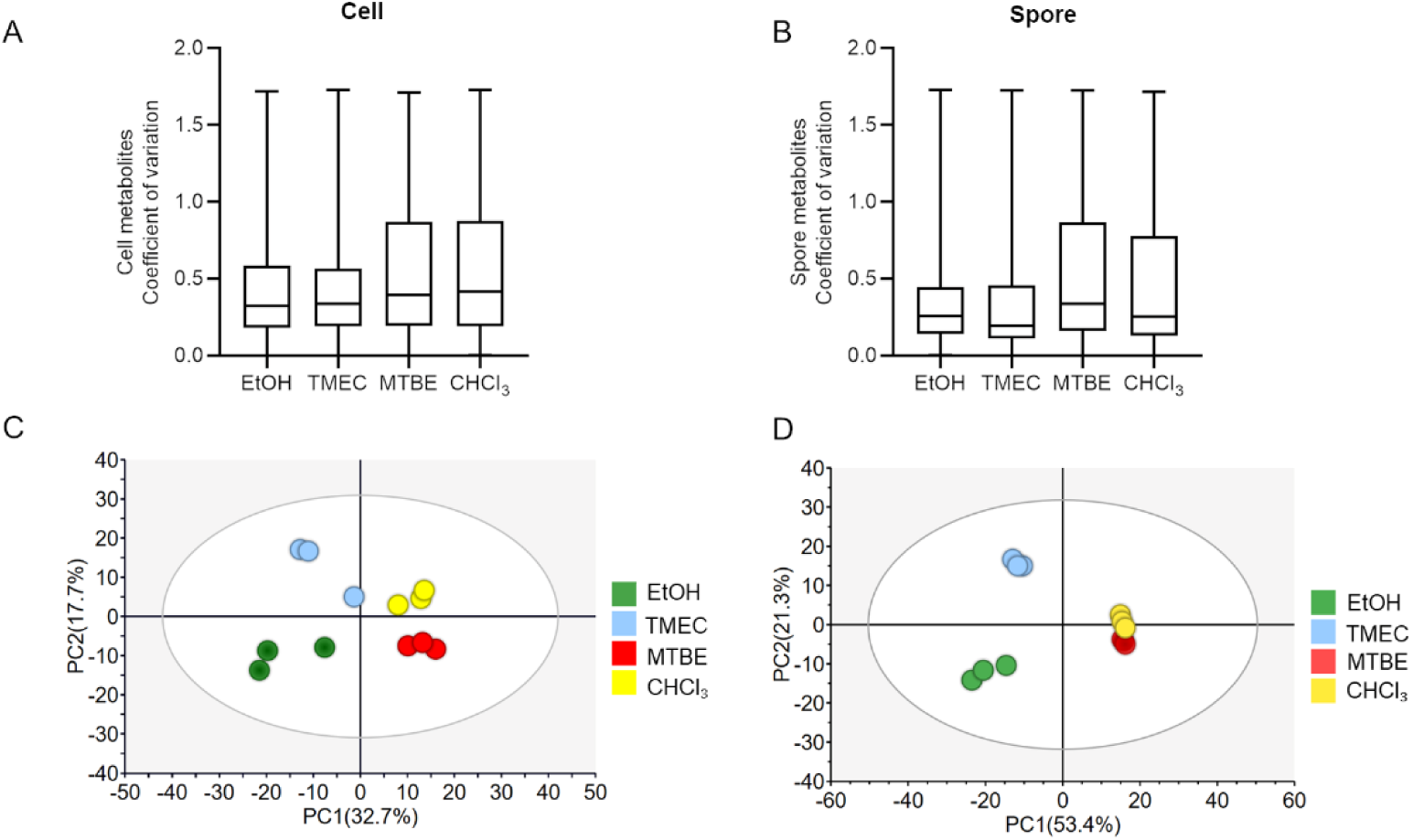
Monophasic methods and biphasic methods showed totally different metabolome profiles. Evaluation of metabolite data quality using different extraction methods for *B. subtilis* cells and spores. A and B, box plots of coefficient of variation (cv) across all the features identified in samples extracted by different methods. C and D, PCA of *B. subtilis* cells and spore’s identified metabolites respectively with different methods.

**Figure S2.**
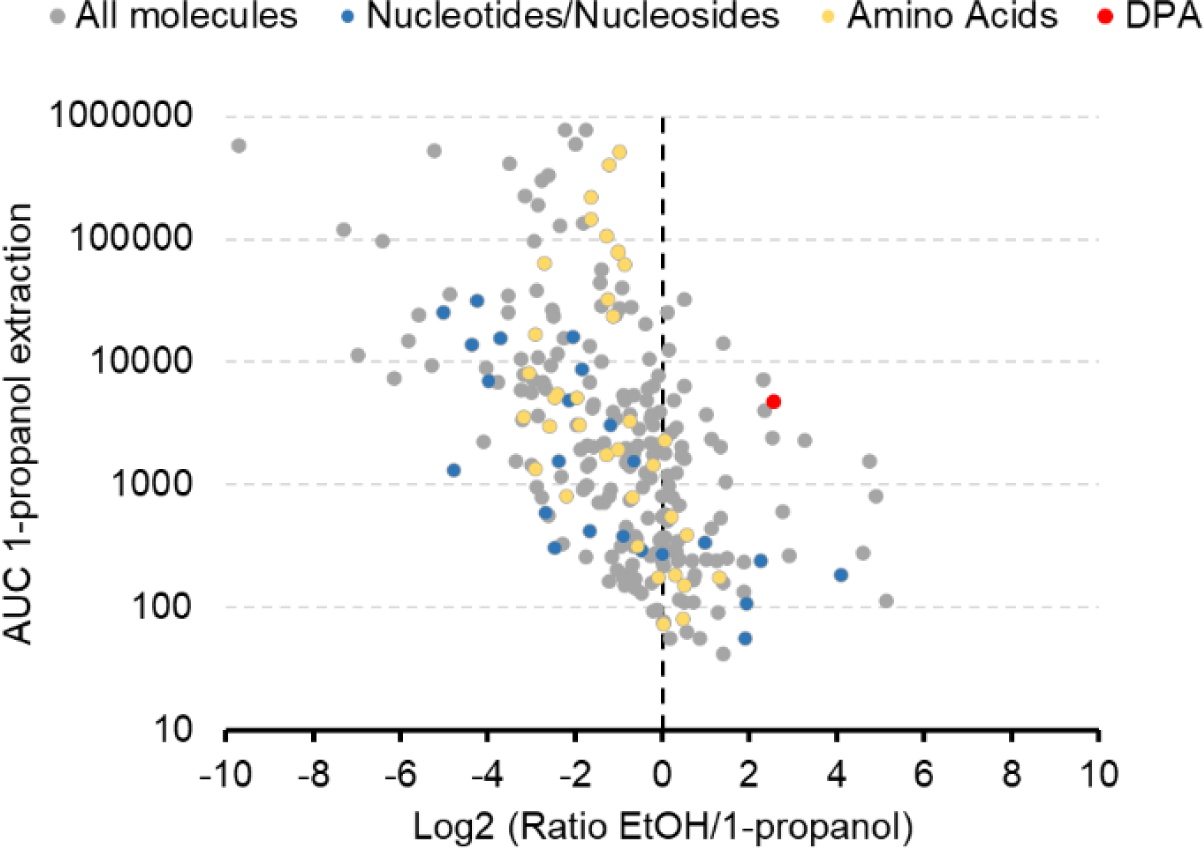
Relative representation of molecules extracted by 60% EtOH with mechanical disruption to boiling 1-propanol extraction in *B. subtilis* spores. X-axis shows ratio between AUC of the two extractions showing molecules with predominant signals in one of the two extractions, Y-axis shows the AUC in the 1-propanol extraction showing the signal intensity of the various molecules in that extraction.

**Figure S3.**
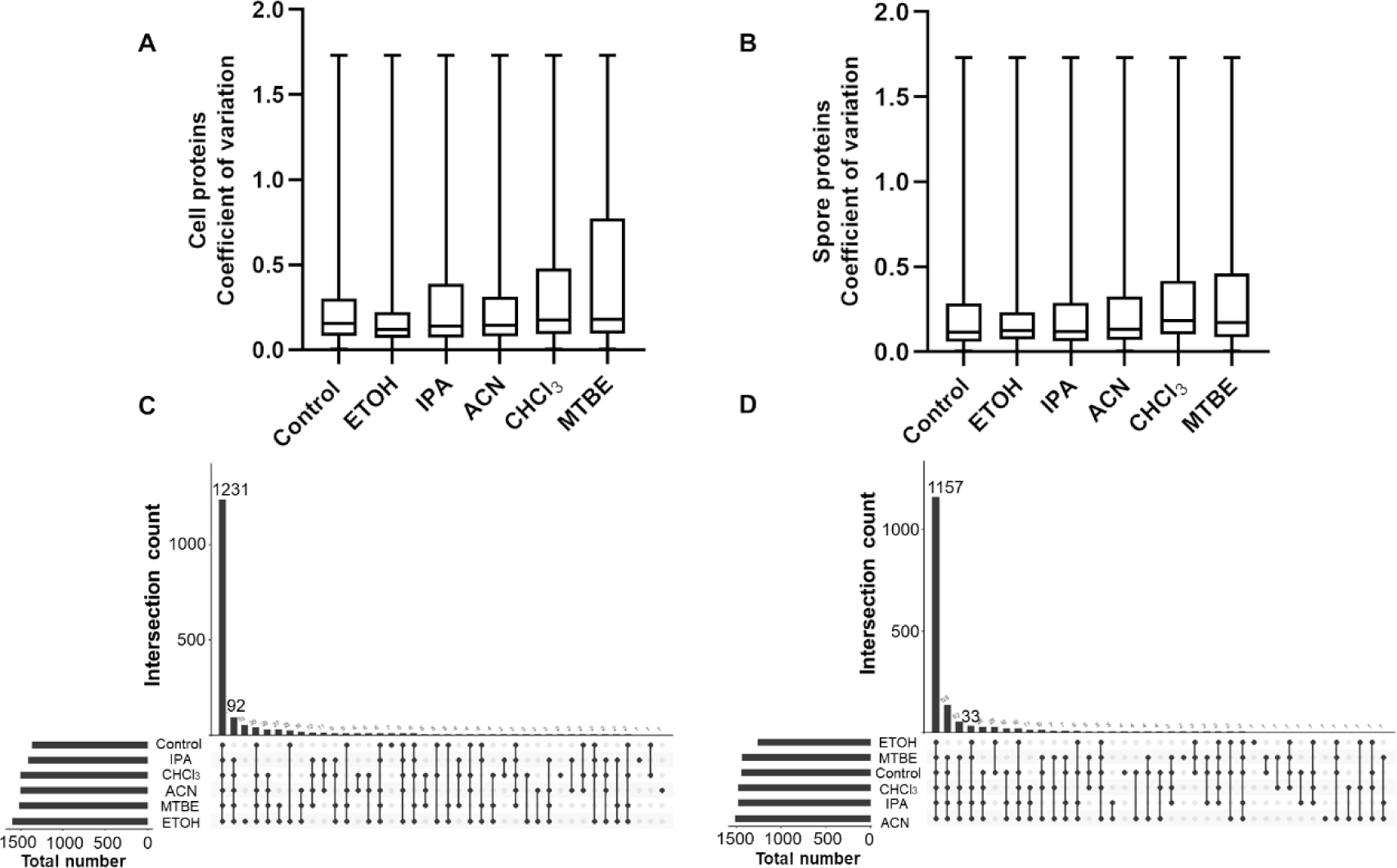
Evaluation of proteome data quality using different extraction methods in *B. subtilis* cells and spores. Control group was lysis by 1% SDS without previous extraction. The A and B, box plots of coefficient of variation (cv) across proteins identified in samples extracted by different methods from *B. subtilis* cells and spores. C and D upset diagram showing the overlapping (shared) and unique metabolites identified in cells and spores by different extraction methods and normal extraction for proteomics.

## Abbreviations

ACN: acetonitrile
MeOH: methanol
IPA: isopropanol
EtOH: ethanol
MTBE: methyl-tert-buthyl-ether
CHCl_3_: chloroform
CV: coefficient of variation
DNA: deoxyribonucleic acid
RNA: ribonucleic acid
EC: energy charge
LC-MS: liquid chromatography–mass spectrometry
LB: Luria-Bertani
MOPS: 3-(nmorpholino) propanesulfonic acid
ABC: ammonium bicarbonate
SDS: sodium-dodecyl sulfate
TCEP: tris(2-carboxyethyl) phosphin hydrochloride
CAA: 2-Chloroacetamide
BCA: bicinchoninic acid
TMEC: combination of two monophasic extractions
SP3: single tube solid phase sample preparation
TIMS: trapped ion mobility spectrometry
MS: mass spectrometer
HILIC: hydrophilic interaction liquid chromatography
FA: formic acid
UHPLC: Ultra high performance liquid chromatography
PASEF: parallel accumulation serial fragmentation
DDA: data dependent acquisition
LFQ: label free quantification
KEGG: kyoto encyclopedia of genes and genomes
GO: gene ontology
PE: phosphatidylethanolamine
PG: phosphatidylglycerol
FA: fatty acyls
LPE: lyso-phosphatidylethanolamine
DAG: diacylglycerol
LPG: lyso-phosphatidylglycerol
PA: phosphatidic acid
TAG: triacylglycerol
PR: prenol lipids
GOCC: gene ontology cell compound.
MF: molecular function
BP: biological process
FDR: false discovery rate.

## References

[1] S.J.C.M. Oomes, A.C.M. van Zuijlen, J.O. Hehenkamp, H. Witsenboer, J.M.B.M. van der Vossen, S. Brul, The characterisation of Bacillus spores occurring in the manufacturing of (low acid) canned products, Int. J. Food Microbiol. 120 (2007) 85–94. https://doi.org/10.1016/j.ijfoodmicro.2007.06.013.

[2] W.L. Nicholson, N. Munakata, G. Horneck, H.J. Melosh, P. Setlow, Resistance of Bacillus Endospores to Extreme Terrestrial and Extraterrestrial Environments, Microbiol. Mol. Biol. Rev. 64 (2000) 548–572.

[3] A.O. Henriques, C.P. Moran Jr., Structure, Assembly, and Function of the Spore Surface Layers, Annu. Rev. Microbiol. 61 (2007) 555–588. https://doi.org/10.1146/annurev.micro.61.080706.093224.

[4] P. Setlow, Spore Resistance Properties, Microbiol. Spectr. 2 (2014) 2.5.11. https://doi.org/10.1128/microbiolspec.TBS-0003-2012.

[5] R.H. Michna, F.M. Commichau, D. Tödter, C.P. Zschiedrich, J. Stülke, SubtiWiki–a database for the model organism Bacillus subtilis that links pathway, interaction and expression information, Nucleic Acids Res. 42 (2014) D692–D698. https://doi.org/10.1093/nar/gkt1002.

[6] K. Nagler, A.O. Krawczyk, A. De Jong, K. Madela, T. Hoffmann, M. Laue, O.P. Kuipers, E. Bremer, R. Moeller, Identification of Differentially Expressed Genes during Bacillus subtilis Spore Outgrowth in High-Salinity Environments Using RNA Sequencing, Front. Microbiol. 7 (2016). https://www.frontiersin.org/articles/10.3389/fmicb.2016.01564 (accessed January 30, 2023).

[7] B. Swarge, W. Abhyankar, M. Jonker, H. Hoefsloot, G. Kramer, P. Setlow, S. Brul, L.J. de Koning, Integrative Analysis of Proteome and Transcriptome Dynamics during Bacillus subtilis Spore Revival, MSphere. 5 (2020) e00463–20. https://doi.org/10.1128/mSphere.00463-20.

[8] Y. Hasin, M. Seldin, A. Lusis, Multi-omics approaches to disease, Genome Biol. 18 (2017) 83. https://doi.org/10.1186/s13059-017-1215-1.

[9] M. Civelek, A.J. Lusis, Systems genetics approaches to understand complex traits, Nat. Rev. Genet. 15 (2014) 34–48. https://doi.org/10.1038/nrg3575.

[10] S. Geng, B.B. Misra, E. de Armas, D.V. Huhman, H.T. Alborn, L.W. Sumner, S. Chen, Jasmonate-mediated stomatal closure under elevated CO2 revealed by time-resolved metabolomics, Plant J. 88 (2016) 947–962. https://doi.org/10.1111/tpj.13296.

[11] C. Buccitelli, M. Selbach, mRNAs, proteins and the emerging principles of gene expression control, Nat. Rev. Genet. 21 (2020) 630–644. https://doi.org/10.1038/s41576-020-0258-4.

[12] S. Donati, M. Kuntz, V. Pahl, N. Farke, D. Beuter, T. Glatter, J.V. Gomes-Filho, L. Randau, C.-Y. Wang, H. Link, Multi-omics Analysis of CRISPRi-Knockdowns Identifies Mechanisms that Buffer Decreases of Enzymes in E. coli Metabolism, Cell Syst. 12 (2021) 56–67.e6. https://doi.org/10.1016/j.cels.2020.10.011.

[13] M. Cascante, S. Marín, Metabolomics and fluxomics approaches, Essays Biochem. 45 (2008) 67–81. https://doi.org/10.1042/BSE0450067.

[14] H.K. Kim, R. Verpoorte, Sample preparation for plant metabolomics, Phytochem. Anal. 21 (2010) 4–13. https://doi.org/10.1002/pca.1188.

[15] M. Salem, M. Bernach, K. Bajdzienko, P. Giavalisco, A Simple Fractionated Extraction Method for the Comprehensive Analysis of Metabolites, Lipids, and Proteins from a Single Sample, J. Vis. Exp. (2017) 55802. https://doi.org/10.3791/55802.

[16] J. Folch, M. Lees, G.H.S. Stanley, A SIMPLE METHOD FOR THE ISOLATION AND PURIFICATION OF TOTAL LIPIDES FROM ANIMAL TISSUES, J. Biol. Chem. 226 (1957) 497–509. https://doi.org/10.1016/S0021-9258(18)64849-5.

[17] V. Matyash, G. Liebisch, T.V. Kurzchalia, A. Shevchenko, D. Schwudke, Lipid extraction by methyl-tert-butyl ether for high-throughput lipidomics, J. Lipid Res. 49 (2008) 1137–1146. https://doi.org/10.1194/jlr.D700041-JLR200.

[18] P. Masson, A.C. Alves, T.M.D. Ebbels, J.K. Nicholson, E.J. Want, Optimization and Evaluation of Metabolite Extraction Protocols for Untargeted Metabolic Profiling of Liver Samples by UPLC-MS, Anal. Chem. 82 (2010) 7779–7786. https://doi.org/10.1021/ac101722e.

[19] J. Kang, L. David, Y. Li, J. Cang, S. Chen, Three-in-One Simultaneous Extraction of Proteins, Metabolites and Lipids for Multi-Omics, Front. Genet. 12 (2021). https://www.frontiersin.org/articles/10.3389/fgene.2021.635971 (accessed January 30, 2023).

[20] C. Coman, F.A. Solari, A. Hentschel, A. Sickmann, R.P. Zahedi, R. Ahrends, Simultaneous Metabolite, Protein, Lipid Extraction (SIMPLEX): A Combinatorial Multimolecular Omics Approach for Systems Biology, Mol. Cell. Proteomics. 15 (2016) 1435–1466. https://doi.org/10.1074/mcp.M115.053702.

[21] H. Meyer, H. Weidmann, M. Lalk, Methodological approaches to help unravel the intracellular metabolome of Bacillus subtilis, Microb. Cell Factories. 12 (2013) 69. https://doi.org/10.1186/1475-2859-12-69.

[22] P. Setlow, A. Kornberg, Biochemical Studies of Bacterial Sporulation and Germination, J. Biol. Chem. 245 (1970) 3637–3644. https://doi.org/10.1016/S0021-9258(18)62974-6.

[23] P. Setlow, A. Kornberg, Biochemical Studies of Bacterial Sporulation and Germination, J. Biol. Chem. 245 (1970) 3645–3652. https://doi.org/10.1016/S0021-9258(18)62975-8.

[24] P. Whittaker, F.S. Fry, S.K. Curtis, S.F. Al-Khaldi, M.M. Mossoba, M.P. Yurawecz, V.C. Dunkel, Use of Fatty Acid Profiles to Identify Food-Borne Bacterial Pathogens and Aerobic Endospore-Forming Bacilli, J. Agric. Food Chem. 53 (2005) 3735–3742. https://doi.org/10.1021/jf040458a.

[25] Y. Song, R. Yang, Z. Guo, M. Zhang, X. Wang, F. Zhou, Distinctness of spore and vegetative cellular fatty acid profiles of some aerobic endospore-forming bacilli, J. Microbiol. Methods. 39 (2000) 225–241. https://doi.org/10.1016/S0167-7012(99)00123-2.

[26] D.L. Nelson, A. Kornberg, Biochemical Studies of Bacterial Sporulation and Germination, J. Biol. Chem. 245 (1970) 1128–1136. https://doi.org/10.1016/S0021-9258(18)63298-3.

[27] P. Setlow, G. Primus, Protein metabolism during germination of Bacillus megaterium spores. I. Protein synthesis and amino acid metabolism, J. Biol. Chem. 250 (1975) 623–630. https://doi.org/10.1016/S0021-9258(19)41942-X.

[28] S. Ghosh, G. Korza, M. Maciejewski, P. Setlow, Analysis of Metabolism in Dormant Spores of Bacillus Species by 31P Nuclear Magnetic Resonance Analysis of Low-Molecular-Weight Compounds, J. Bacteriol. 197 (2015) 992–1001. https://doi.org/10.1128/JB.02520-14.

[29] D. Li, T.V. Truong, T.M. Bills, B.C. Holt, D.N. VanDerwerken, J.R. Williams, A. Acharya, R.A. Robison, H.D. Tolley, M.L. Lee, GC/MS Method for Positive Detection of Bacillus anthracis Endospores, Anal. Chem. 84 (2012) 1637–1644. https://doi.org/10.1021/ac202606x.

[30] G. Korza, M. Goulet, A. DeMarco, J. Wicander, P. Setlow, Role of Bacillus subtilis Spore Core Water Content and pH in the Accumulation and Utilization of Spores’ Large 3-Phosphoglyceric Acid Depot, and the Crucial Role of This Depot in Generating ATP Early during Spore Germination, Microorganisms. 11 (2023) 195. https://doi.org/10.3390/microorganisms11010195.

[31] G. Bertani, Studies on lysogenesis i, J. Bacteriol. 62 (1951) 293–300. https://doi.org/10.1128/jb.62.3.293-300.1951.

[32] R. Kort, A.C. O’Brien, I.H.M. van Stokkum, S.J.C.M. Oomes, W. Crielaard, K.J. Hellingwerf, S. Brul, Assessment of Heat Resistance of Bacterial Spores from Food Product Isolates by Fluorescence Monitoring of Dipicolinic Acid Release, Appl. Environ. Microbiol. 71 (2005) 3556– 3564. https://doi.org/10.1128/AEM.71.7.3556-3564.2005.

[33] H. Meyer, M. Liebeke, M. Lalk, A protocol for the investigation of the intracellular Staphylococcus aureus metabolome, Anal. Biochem. 401 (2010) 250–259. https://doi.org/10.1016/j.ab.2010.03.003.

[34] A.D. Southam, L.D. Haglington, L. Najdekr, A. Jankevics, R.J.M. Weber, W.B. Dunn, Assessment of human plasma and urine sample preparation for reproducible and high-throughput UHPLC-MS clinical metabolic phenotyping, Analyst. 145 (2020) 6511–6523. https://doi.org/10.1039/D0AN01319F.

[35] E.G. Bligh, W.J. Dyer, A RAPID METHOD OF TOTAL LIPID EXTRACTION AND PURIFICATION, (n.d.).

[36] B.N. Swarge, W. Roseboom, L. Zheng, W.R. Abhyankar, S. Brul, C.G. de Koster, L.J. de Koning, “One-Pot” Sample Processing Method for Proteome-Wide Analysis of Microbial Cells and Spores, Proteomics Clin. Appl. 12 (2018) e1700169. https://doi.org/10.1002/prca.201700169.

[37] L. Zheng, W. Abhyankar, N. Ouwerling, H.L. Dekker, H. van Veen, N.N. van der Wel, W. Roseboom, L.J. de Koning, S. Brul, C.G. de Koster, Bacillus subtilis Spore Inner Membrane Proteome, J. Proteome Res. 15 (2016) 585–594. https://doi.org/10.1021/acs.jproteome.5b00976.

[38] X. Gao, B.N. Swarge, H.L. Dekker, W. Roseboom, S. Brul, G. Kramer, The Membrane Proteome of Spores and Vegetative Cells of the Food-Borne Pathogen Bacillus cereus, Int. J. Mol. Sci. 22 (2021) 12475. https://doi.org/10.3390/ijms222212475.

[39] A. Tanca, A. Palomba, S. Pisanu, M. Deligios, C. Fraumene, V. Manghina, D. Pagnozzi, M.F. Addis, S. Uzzau, A straightforward and efficient analytical pipeline for metaproteome characterization, Microbiome. 2 (2014) 49. https://doi.org/10.1186/s40168-014-0049-2.

[40] K. Hayoun, D. Gouveia, L. Grenga, O. Pible, J. Armengaud, B. Alpha-Bazin, Evaluation of Sample Preparation Methods for Fast Proteotyping of Microorganisms by Tandem Mass Spectrometry, Front. Microbiol. 10 (2019) 1985. https://doi.org/10.3389/fmicb.2019.01985.

[41] C.S. Hughes, S. Moggridge, T. Müller, P.H. Sorensen, G.B. Morin, J. Krijgsveld, Single-pot, solid-phase-enhanced sample preparation for proteomics experiments, Nat. Protoc. 14 (2019) 68–85. https://doi.org/10.1038/s41596-018-0082-x.

[42] J. Edwards-Hicks, M. Mitterer, E.L. Pearce, J.M. Buescher, Metabolic Dynamics of In Vitro CD8+ T Cell Activation, Metabolites. 11 (2020) 12. https://doi.org/10.3390/metabo11010012.

[43] J.H. Martin, Fatty acids of vegetative cells and spores of Bacillus licheniformis, J. Dairy Sci. 59 (1976) 1830–1834. https://doi.org/10.3168/jds.s0022-0302(76)84444-x.

[44] I.R. León, V. Schwämmle, O.N. Jensen, R.R. Sprenger, Quantitative assessment of in-solution digestion efficiency identifies optimal protocols for unbiased protein analysis, Mol. Cell. Proteomics MCP. 12 (2013) 2992–3005. https://doi.org/10.1074/mcp.M112.025585.

[45] S. Englard, S. Seifter, [22] Precipitation techniques, in: M.P. Deutscher (Ed.), Methods Enzymol., Academic Press, 1990: pp. 285–300. https://doi.org/10.1016/0076-6879(90)82024-V.

[46] A.A. Green, W.L. Hughes, [10] Protein fractionation on the basis of solubility in aqueous solutions of salts and organic solvents, 1 (1955) 67–90. https://doi.org/10.1016/0076-6879(55)01014-8.

[47] P. Setlow, Spores of Bacillus subtilis: their resistance to and killing by radiation, heat and chemicals, J. Appl. Microbiol. 101 (2006) 514–525. https://doi.org/10.1111/j.1365-2672.2005.02736.x.

[48] Ž. Kolenc, N. Pirih, P. Gretic, T. Kunej, Top Trends in Multiomics Research: Evaluation of 52 Published Studies and New Ways of Thinking Terminology and Visual Displays, OMICS J. Integr. Biol. 25 (2021) 681–692. https://doi.org/10.1089/omi.2021.0160.

[49] F. Zheng, X. Zhao, Z. Zeng, L. Wang, W. Lv, Q. Wang, G. Xu, Development of a plasma pseudotargeted metabolomics method based on ultra-high-performance liquid chromatography-mass spectrometry, Nat. Protoc. 15 (2020) 2519–2537. https://doi.org/10.1038/s41596-020-0341-5.

[50] L. Brennan, P. Owende, Biofuels from microalgae--A review of technologies for production, processing, and extractions of biofuels and co-products, Renew. Sustain. Energy Rev. 14 (2010) 557–577.

[51] T. Dong, E.P. Knoshaug, P.T. Pienkos, L.M.L. Laurens, Lipid recovery from wet oleaginous microbial biomass for biofuel production: A critical review, Appl. Energy. 177 (2016) 879–895. https://doi.org/10.1016/j.apenergy.2016.06.002.

[52] H. Sati, M. Mitra, S. Mishra, P. Baredar, Microalgal lipid extraction strategies for biodiesel production: A review, Algal Res. 38 (2019) 101413. https://doi.org/10.1016/j.algal.2019.101413.

[53] C.J. Seneviratne, T. Suriyanarayanan, A.S. Widyarman, L.S. Lee, M. Lau, J. Ching, C. Delaney, G. Ramage, Multi-omics tools for studying microbial biofilms: current perspectives and future directions, Crit. Rev. Microbiol. 46 (2020) 759–778. https://doi.org/10.1080/1040841X.2020.1828817.

[54] L. Man, W.P. Klare, A.L. Dale, J.A. Cain, S.J. Cordwell, Integrated mass spectrometry-based multi-omics for elucidating mechanisms of bacterial virulence, Biochem. Soc. Trans. 49 (2021) 1905–1926. https://doi.org/10.1042/BST20191088.

[55] C. Kerepesi, J.E. Szabó, V. Papp-Kádár, O. Dobay, D. Szabó, V. Grolmusz, B.G. Vértessy, Life without dUTPase, Front. Microbiol. 7 (2016). https://www.frontiersin.org/articles/10.3389/fmicb.2016.01768 (accessed January 30, 2023).

[56] N. Kamatani, H.A. Jinnah, R.C.M. Hennekam, A.B.. P. van Kuilenburg, Purine and Pyrimidine Metabolism, in: Ref. Module Biomed. Sci., Elsevier, 2014. https://doi.org/10.1016/B978-0-12-801238-3.05567-7.

[57] A.R. Bate, R. Bonneau, P. Eichenberger, Bacillus subtilis Systems Biology: Applications of -Omics Techniques to the Study of Endospore Formation, Microbiol. Spectr. 2 (2014). https://doi.org/10.1128/microbiolspec.TBS-0019-2013.

